# The clinical and molecular significance associated with STING signaling in estrogen receptor-positive early breast cancer

**DOI:** 10.1101/2020.07.23.217398

**Authors:** EE Parkes, MP Humphries, E Gilmore, FA Sidi, V Bingham, SM Phyu, SG Craig, C Graham, J Miller, D Griffin, RD Kennedy, SF Bakhoum, S McQuaid, M Salto-Tellez, NE Buckley

## Abstract

STING signaling in cancer is a crucial component of response to immunotherapy and other anti-cancer treatments. Conversely, STING signaling can promote tumor invasion and metastasis. Currently, there is no robust method of measuring STING activation in cancer. Here, we describe an immunohistochemistry-based assay with digital pathology assessment of STING in tumor cells. Using this novel approach, we identify perinuclear-localized expression of STING (pnSTING) in estrogen receptor-positive (ER+) breast cancer as an independent predictor of good prognosis, associated with immune cell infiltration and upregulation of immune checkpoints. Tumors with low pnSTING are immunosuppressed with increased infiltration of “M2” -polarised macrophages. In ER-disease, pnSTING does not have a significant prognostic role, and STING appears to be uncoupled from interferon responses. Importantly, a gene signature defining low pnSTING expression in ER+ disease is predictive of poor prognosis in independent datasets. Low pnSTING is associated with chromosomal instability, *MYC* amplification and mTOR signaling, suggesting novel therapeutic approaches for this subgroup.

## Introduction

Avoiding immune destruction and tumor-promoting inflammation are immune hallmarks of cancer, with the innate immune cyclic GMP-AMP synthase (cGAS) - STimulator of INterferon Genes (STING) pathway involved in both ^1–3^. The cGAS-STING pathway is a focal point of innate immune responses, activated when cGAS detects cytosolic DNA, producing 2’3’cGAMP and resulting in subsequent stimulation of STING and downstream TBK1-IRF3 and NFκB-RelB pathways ^4^.

Activation of STING in response to DNA damaging therapies such as ionizing radiation has been reported as essential for a direct antitumor and abscopal response, as well as implicated in subsequent inflammation-mediated radioresistance. ^5–7^. Activation of the STING pathway via cGAS has been identified as a prerequisite for response to immune checkpoint blockade (ICB) ^8^. STING agonists, where delivery of intratumoral cyclic dinucleotides activate STING responses within the tumor microenvironment, (TME) have demonstrated synergy with ICB in preclinical models ^9,10^. Trials are ongoing to determine the safety and efficacy of STING agonists in combination with ICB in advanced metastatic cancer ^11,12^.

We previously identified a STING-driven chemokine response signature in DNA repair-deficient breast cancers, which was associated with improved outcome in the context of standard-of-care chemotherapy ^13^. However, chronic stimulation of cGAS-STING via micronuclei in chromosomally unstable cancers results in inactivation of interferons and instead promotes downstream NFκB-RelB-mediated metastasis ^3^. The intrinsic STING activation or suppression inherent in cancer is a therefore a potential predictive biomarker for immune or other therapies. Furthermore, identifying immune and gene expression correlates with STING activation could suggest rational combination approaches in STING-agonist resistant cancers. Expression of *TMEM173*, encoding STING, moderately correlates with immune infiltration but only poorly correlates with expression of downstream components TBK1 and IRF3, suggesting that *TMEM173* expression may be a weak indicator of STING activity ^14^. A specific STING activation assay would be a valuable tool in addressing these important questions. However, given the diversity of downstream chemokine responses, transient nature of TBK1-IRF3 activation and crosstalk with other nucleic acid sensing pathways, such an approach has been challenging to develop.

## Materials and Methods

### Cell lines

MDA-MB-436-EV (described in ^13^) were maintained in 50% Leibovitz’s L-15 / 50% RPMI (ThermoFisher Scientific, Paisley, UK) supplemented with 10% fetal bovine serum. Cells were confirmed as mycoplasma free in routine lab testing.

### Immunofluorescence

Cells were seeded on a glass coverslip in 6-well plates and incubated overnight at 37 °C. Cells were then transfected with 10 μg/ml 2’3’cGAMP (Invivogen, Toulouse, France). After 1 hour, medium was removed and cells fixed in methanol at −20 °C for 20 minutes. Slides were covered with a solution of 0.5% (v/v) TritonX-100 in PBS and incubated at room temperature for 15 minutes. Following 1 hour of blocking in 5% (v/v) FBS with 0.2% (v/v) Tween20 in PBS, anti-STING antibody (Cell Signaling Technology, Leiden, Netherlands) was added at 1:600 and slides incubated at 4 °C overnight. Fluorescently labelled antibody (anti-rabbit IgG, Alexa Fluor488-conjugated, Life Technologies) was added at 1:1500. Following 3 hours incubation at room temperature, DAPI was added for 15 minutes. Slides were mounted using ProLong Gold Antifade Mountant (ThermoFisher Scientific).

### Breast TMA

The cohort of 300 de-novo breast cancer patients, their clinical, pathological and outcome parameters and the construction of the tissue microarrays (TMAs) used in the present study has been previously described ^15–17^. Cases were diagnosed in Northern Ireland from 1997 – 2009 with ethical approval granted by the Northern Ireland Biobank ^18^ (NIB12-0017, NIB15-0168). STING IHC was approved under NIB19-301.

### Immunohistochemistry

All immunohistochemistry (IHC) was performed in a hybrid laboratory (Precision Medicine Centre of Excellence), awarded UK Clinical Pathology Accreditation. Sections were cut from the TMA blocks for H&E staining and IHC for the range of biomarkers described in **Suppl. Table 1**. Briefly, sections for IHC were cut at 4 microns on a rotary microtome, dried at 37 °C overnight, with IHC then performed on automated immunostainers (Leica Bond-Max, Milton Keynes, UK or Ventana BenchMark, Tucson, AZ). Each biomarker was initially validated on carefully selected control tissues. Antigen-binding sites were detected with a polymer-based detection system (Leica Biosystems UK, Cat. No. DS9800 or Ventana USA Cat. No. 760-700 and Cat. No. 760-500). All sections were subsequently visualized with diaminobenzidine (DAB), counterstained with hematoxylin, and tape mounted using a Sakura Autostainer (Sakura Finetek Europe, Rijn, Netherlands). All slides were scanned on an Aperio AT2 digital scanner (Leica Biosystems, Milton Keynes, UK) at x 40 magnification.

For multiplexing immunofluorescence, 4 micron sections were obtained from cases observed to demonstrate perinuclear STING by DAB IHC. Sections were stained with validated methods, as described ^19,20^. Staining was performed on a Leica Bond-Max (Leica Biosystems UK, Milton Keynes), using Opal 4-Color Automation IHC Kit (CK/STING/DAPI) (Cat. No. NEL820001KT, Akoya Biosciences, Marlborough, MA). According to the manufacturer’s instructions, all retrieval methods and staining steps for Opal were optimized and are detailed in **Suppl. Table 1**. All multiplex slides were scanned on a Vectra Polaris (Akoya Biosciences) at ×20.

### Image Analysis

Digital pathological analysis of all IHC stained TMA slides was performed using QuPath, an open-source image analysis program ^21^. All x 40 scanned slides were imported, dearrayed, and tissue detection carried out to identify the areas of tissue available for cellular analysis. Cores were removed following strict exclusion criteria; e.g., tissue cores that contained < 100 tumor cells were removed from analysis. Rigorous QC steps were taken to remove necrosis, tissue folds, and entrapped normal structures; this was confirmed by a second reviewer with frequent consultation. For each biomarker, positive staining was defined as the presence of any discernible DAB positivity localized in either the nucleus, membrane and/or cytoplasm depending on the known biological expression. After intensive quality control Intensity thresholds were set for cellular DAB detection. Percentage positive data was extracted from each TMA core and averaged across replicates.

### Gene expression analysis

As described previously^15^, total RNA was extracted from macrodissected formalin-fixed paraffin embedded (FFPE) tumor samples using the Roche High Pure RNA Paraffin Kit. Following amplification using the NuGEN WT-Ovation FFPE System (NuGEN, San Carlos, CA), total RNA was hybridized to the Almac Breast Cancer DSA (Affymetrix, Santa Clara, CA).

### Statistical analysis

Correlation was carried out using Spearman test in the context of continuous data or Mann Whitney and Kruskall-Wallis test as appropriate in the context of categorical data. Fishers exact or Chi squared analysis were carried out as appropriate using Graphpad Prism v8 Software. Data was transformed for graphing purposes only.

Heatmaps were generated using robust z-score transformed data. Survival analysis was carried out using GraphPad Prism v8 with log rank hazard ratios and p-values reported unless stated otherwise. Multivariate analysis was carried out using the R package “survival” in the context of Age (<40, 40-49, 50-59, >60), T stage (1, 2, 3, 4) and lymph node status (positive or negative).

### Gene Set Enrichment Analysis

Gene Set Enrichment Analysis (GSEA) was carried out using the publicly available GSEA tool (v4.0.3) (www.gsea-msigdb.org) using the Hallmarks gene sets (h.all.v7.1)

### STING activity gene signature

GSEA was used to identify the top 25 ranked genes upregulated in the ER+ cases called “STING Perinuclear Low” by IHC analysis. The negative sum of the expression of the 25 genes was calculated from gene expression microarray datasets using gene-level summarizing where appropriate.

### External datasets

Datasets were selected based on the following criteria: Availability of overall and/or progression-free survival data, availability of ER status measured by IHC, availability of microarray mRNA expression data.

The METABRIC ^22^ and TCGA Nature 2012 cohorts ^23^ were accessed via the cBioPortal website ^24,25^. From METABRIC, the 1338 cases for analysis were selected based on “Breast Invasive Ductal Carcinoma”, availability of ER status measured by IHC and availability of data describing chemotherapy status. For the TCGA Nature 2012 cohort, the 519 cases for analysis were selected based on availability of ER status and mRNA expression data. The Wang et al. dataset ^26^ was accessed via GEO datasets (GSE2034) with the associated clinical information used to select ER+ cases.

STING gene expression scores were generated and high vs low (above or below median) were compared using available genomic and clinical information.

## Results

### Perinuclear STING expression predicts prognosis in ER positive breast cancer

To address the role of STING activation and relationship with the tumor immune microenvironment, we utilized a previously described early stage breast tumor TMA ^15,16^. Following review for quality control, STING IHC was available for 156 tumors (**Suppl. Table 2**). We first assessed the expression of STING in a subset of tumor cores. Four distinct patterns of STING expression were observed (**Fig. 1a-d, Suppl. Fig. 1a-b**), where STING expression was high throughout the tumor, high in stromal or tumor epithelial compartment alone, or universally low/absent. In tumors with epithelial cell STING expression, we noted a proportion of cells with a distinctive perinuclear pattern of STING staining. As STING migrates to perinuclear microsomes when activated ^27,28^ (**Suppl. Fig. 1c**), we sought to measure perinuclear STING expression as representative of tumor cell STING activity. Therefore, we applied a digital pathology workflow (**Fig. 1e**) to calculate perinuclear STING (pnSTING) expression in the DAB IHC digital images. Within STING positive tumors, we were able to determine tumors with low (**Fig. 1f**) or high proportions of STING active cells (**Fig. 1g**).

**Figure 1:**
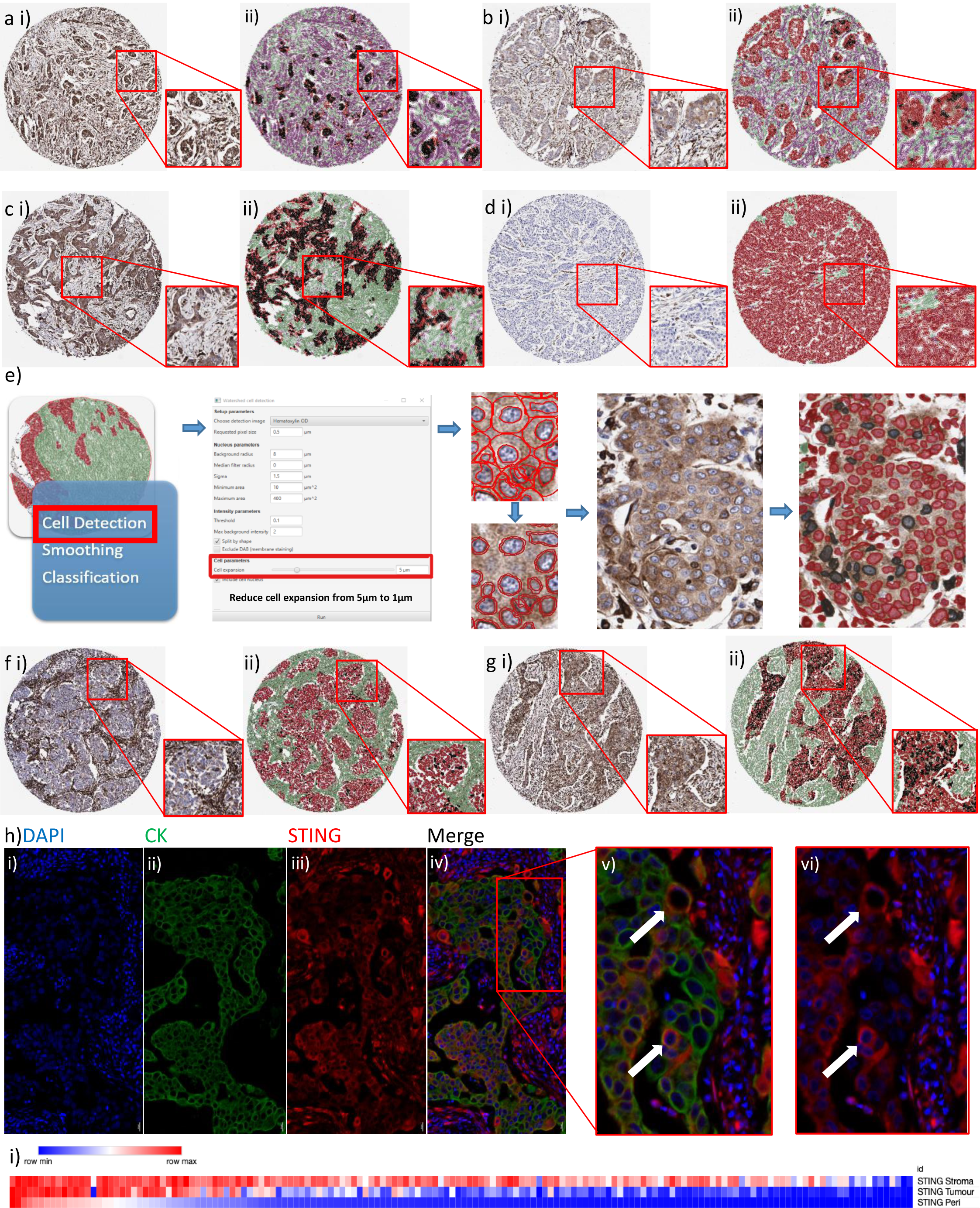
STING immunohistochemistry in breast cancer. Immunohistochemistry (IHC) images representing **a**) high expression of STING in both tumor and stromal compartments, **b**) low expression of STING in tumor with high expression in stroma, **c**) high expression of STING in tumor with low expression in stroma and **d**) low expression of STING in both tumor and stromal compartments. Magnification: cores x 4, inset x 8. **e**) QuPath workflow for perinuclear STING analysis. Black = perinuclear STING positive cells, Red = perinuclear STING negative cells. IHC images representing **f**) low but detectable perinuclear STING in an otherwise STING positive tumor and **g**) high perinuclear STING expression. Magnification: cores x 4, inset x 8. **h**) Multiplex IHC of tumor section with DAPI, STING (red) and cytokeratin (CK, green). Co-localization of STING and CK is demonstrated in in **v**) (with CK) and **vi**) (without CK), indicated by the white arrows. Magnification: **i-iv**) x 10, **v-vi**) x 20. **i**) Correlation of stromal, tumor and perinuclear STING (absolute scores measured as percentage of positive cells) in breast cancer IHC cases. Stromal v. tumor: R = 0.7240, p < 0.0001, Stromal v. perinuclear: R = 0.6916, p < 0.0001, Tumor v. perinuclear: R = 0.8496, p < 0.0001.

To investigate whether observed pnSTING staining was found in tumor epithelial cells or infiltrating immune cells, we performed multiplex studies on selected sections. Close examination verified that pnSTING staining cells were within cytokeratin (CK) positive tumor nests, confirming these cells were not of immune origin (**Fig. 1h**). Nearly all cases had some level of STING expression within the stroma, with over the majority of these also expressing STING within tumor cells (R = 0.6916, p < 0.0001). While total STING levels within the tumor closely correlated with high pnSTING positive cells (R = 0.850, p < 0.0001), pnSTING high tumors were a clearly defined subset within STING-expressing tumors (**Fig. 1i, Suppl. Fig. 1d**).

By measuring the percentage of positive pnSTING cells within the tumor, as delineated by a machine learning digital pathology classifier, we found that breast tumors with high pnSTING (pnSTING^high^, >median) predicted significantly improved relapse free survival (RFS) (HR = 0.405, 95% CI 0.210-0.778, p = 0.0096) compared to either stromal or whole cell tumor STING expression alone (HR = 0.553, 95% CI 0.29-1.05, p = 0.090; HR = 0.554, 95% CI = 0.291-1.06, p = 0. 077 respectively) (**Fig. 2ai-iii, Table 1**). However, the predictive ability of pnSTING expression was independently significant in the ER + population (multivariate analysis HR = 0.206, 95% CI 0.059-0.727, p = 0.014), and did not predict RFS in ER - disease (HR = 0.810, 95% CI 0.309-2.12, p = 0.663) (**Fig. 2bi-ii, Table 2**). While overall survival was numerically improved in the ER + pnSTING^high^ subgroup, this was not significant at 60 months (HR = 0.6288, 95% 0.2201-1.796) (**Suppl. Fig. 2a-b**). Using consensus breast subgroups^29^ pnSTING^high^ predicted RFS in the Luminal A (ER+, HER2-, Ki67-) subgroup (HR = 0.130, 95% CI = 0.002 – 0.757, p = 0.0232) (**Suppl. Fig. 2c-d, Suppl. Table 3**). Comparing pnSTING^high^ and pnSTING^low^ ER+ cases, no significant difference in PAM50 subtype, St. Gallen subtype, grade, age, T or N status or treatment received was observed (**Fig. 2c-d, Suppl. Fig. 2e**).

**Table 1:** STING expression and relapse free survival in discovery cohort (ER+ and ER- breast cancer)

| **Relapse Free Survival** |  | **N(n)** | **HR** | **95% CI** | **p-value** |
| --- | --- | --- | --- | --- | --- |
|  |  | 156 (37) | Univariate |  |  |
| **STING Stroma** | low | 78 (23) | 1 |  |  |
|  | high | 78 (14) | 0.5528 | 0.29-1.054 | 0.0899 |
| **STING Tumor** | low | 78 (23) | 1 |  |  |
|  | high | 78 (14) | 0.5543 | 0.2908-1.057 | 0.0773 |
| **STING Perinuclear** | low | 78 (26) | 1 |  |  |
|  | high | 78 (11) | 0.4046 | 0.2104-0.778 | 0.0096** |

HR: Hazard ratio, CI: Confidence interval; n: events

**Table 2:** STING expression and relapse free survival in ER+ and ER- breast cancer.

| **ER+** |  | **N(n)** | **HR** | **95% CI** | **p-value** |
| --- | --- | --- | --- | --- | --- |
| **Relapse Free Survival** |  | 92 (19) | Univariate |  |  |
| **STING Perinuclear** | low | 50 (16) | 1 |  |  |
|  | high | 42 (3) | 0.1955 | 0.0795 – 0.4805 | 0.0038** |
| **Multivariate** |  |  |  |  |  |
| **STING Perinuclear** | low | 50 (16) | 1 |  |  |
|  | high | 42 (3) | 0.2067 | 0.0588-0.7265 | 0.014* |
| **ER-** |  |  |  |  |  |
| **Relapse Free Survival** |  | 64 (17) | Univariate |  |  |
| **STING Perinuclear** | low | 27 (8) | 1 |  |  |
|  | high | 36 (9) | 0.8098 | 0.3089-2.123 | 0.6633 |

Multivariate in context of age (<40, 40-49, 50-59, >60), T status (1, 2, 3,4) and Lymph node (pos/neg). No other factor was significant.

**Figure 2:**
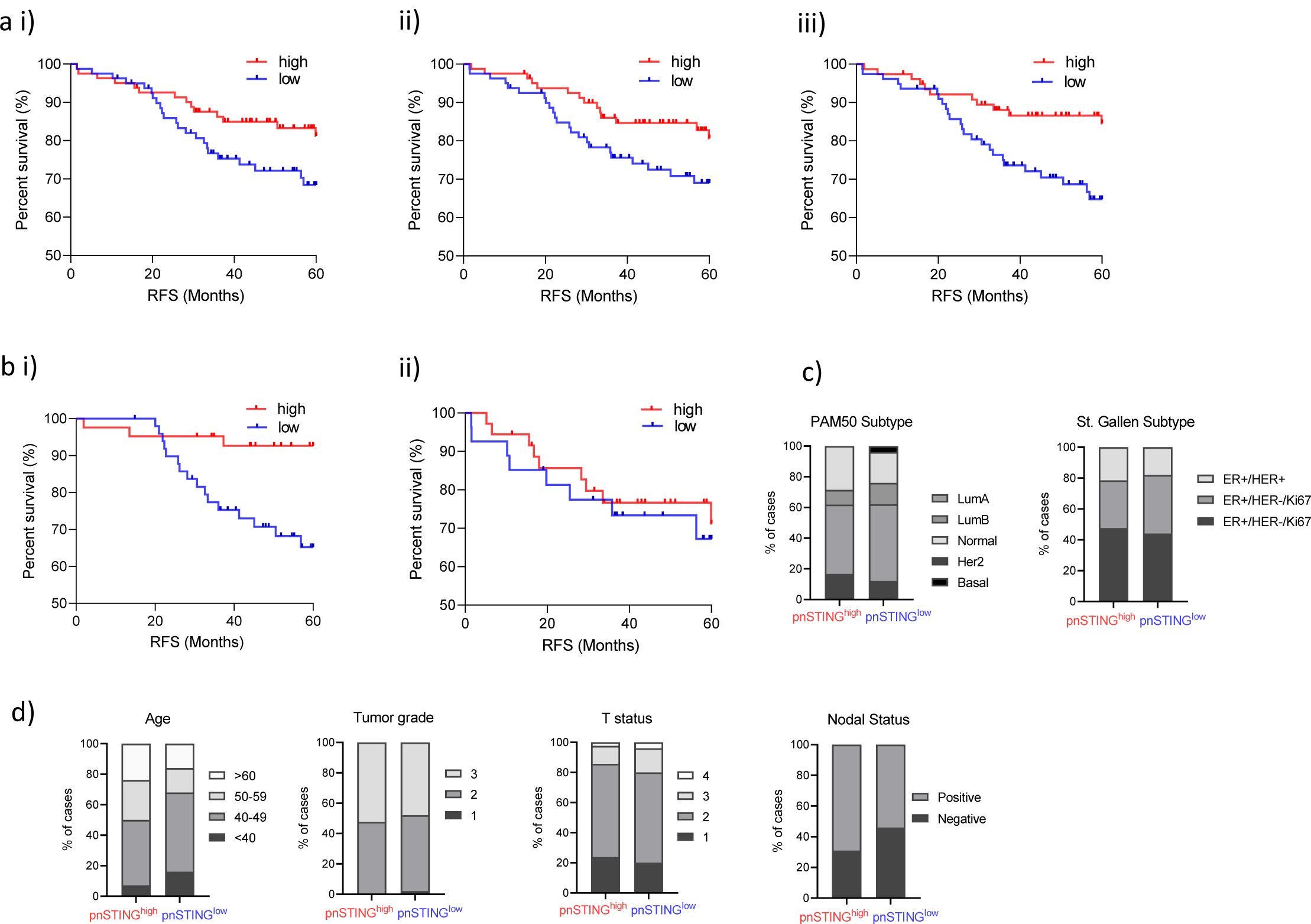
pnSTING IHC score predicts outcome in ER+ breast cancer. **a**) Kaplan-Meier Curve of relapse free survival (RFS) stratified based on high (above median) or low (below median) of STING expression in the **i**) stromal compartment, **ii**) tumour epithelial compartment or **iii**) perinuclear region. **b**) Kaplan Meier Curve of relapse free survival (RFS) stratified based on high (above median of all cases) or low (below median of all cases) of STING expression in the perinuclear region in **i**) ER positive (ER+) and **ii**) ER negative (ER-) cases. **c**) Stacked bar chart of the percentage of ER+ patients stratified based on high (above median) or low (below median) of STING expression in the perinuclear region based PAM50 subtype and St. Gallen subtype. **d**) Stacked bar chart of the percentage of ER+ patients stratified based on high or low pnSTING expression detailing clinicopathological characteristics of Age, Tumor Grade, T stage, N status (from left to right).

### Perinuclear STING expression correlates with the immune landscape of breast cancer

Using matched IHC and array-based gene expression profiling (GEP) we characterized the immune landscape in relation to pnSTING expression. Using digital pathological analysis, we measured T- and B-cell markers, innate immune populations and immune checkpoint expression within tumor and surrounding stroma. In ER+ cases, a significant association of CD3^+^, CD4^+^, CD8^+^ and CD45RO^+^ cells was identified with pnSTING^high^ tumors (**Fig. 3a-b, Suppl. Fig. 3a**). In addition, expression of immune checkpoints PD-L1 (SP263 and SP142), IDO1 and ICOS significantly correlated with pnSTING^high^ tumors. In ER-cases, pnSTING did not significantly correlate with T cell markers except CD3 stromal expression, suggesting an uncoupling of tumor cell STING activity and immune responses in ER-breast cancer. However, a significant increase in CD68^+^ macrophages was noted in ER-pnSTING^high^ tumors compared to pnSTING^low^ (p = 0.008, **Suppl. Fig. 3b**). This trend was not observed in ER+ tumors.

**Figure 3:**
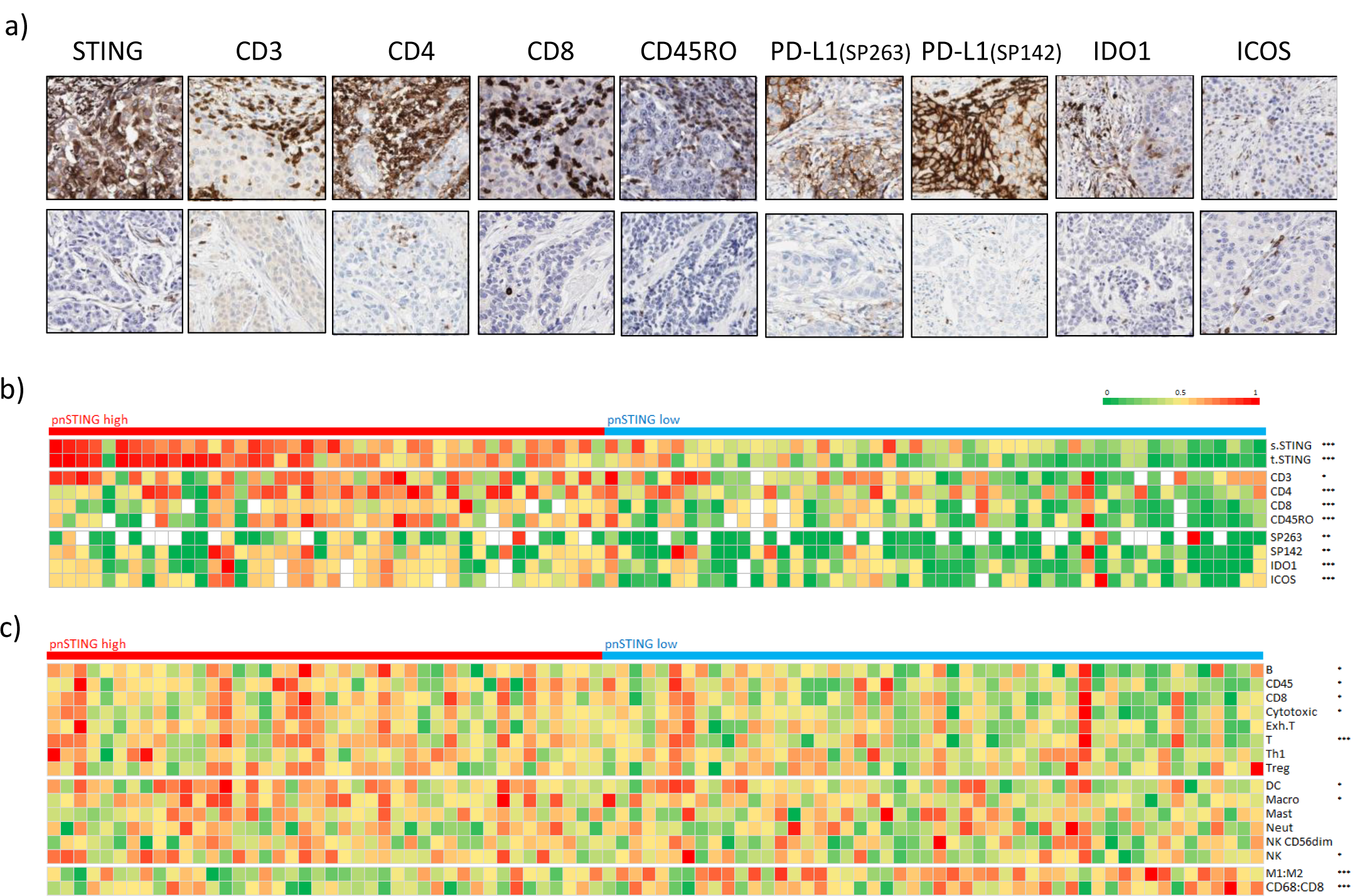
pnSTING and immune correlates in ER+ breast cancer. **a**) Representative IHC images showing high (top panel) and low (lower panel) expression of perinuclear STING, CD3, CD4, CD8, CD45RO, PD-L1 (SP263), PD-L1 (SP142), IDO1 and ICOS. **b**) Heatmap of normalized expression measured by IHC of Perinuclear STING, stroma STING, tumor STING, CD3, CD4, CD8, CD45RO, PD-L1 measured by SP263 and SP142, IDO1 and ICOS in ER+ breast cancer cases. **c**) Heatmap of normalized immune scores derived from deconvolution of microarray data in ER+ breast cancer cases. Correlation between markers and pnSTING stratified based on high (above median) and low (below median) was assessed using the Krushall Wallis test on non-transformed data with *, ** and *** indicating a p values of < 0.05, < 0.01 and < 0.001 respectively.

Using gene expression signatures for predicted immune cell infiltration ^30^, we confirmed significant association of similar lymphocytic populations in the TME to that identified on IHC analysis (**Fig. 3c**). Additionally, dendritic cell and natural killer (NK) cell infiltration were found to be significantly associated with pnSTING^high^ tumors, in keeping with dendritic cells as a key source of PD-L1 and in promoting T-cell responses ^31^. Importantly, gene expression of STING did not strongly correlate with pnSTING scores in ER+ tumors (**Fig. 4a**). No significant associations between predicted immune populations from GEP and pnSTING expression were identified in ER-tumors (**Suppl. Fig. 3c**). However, in ER+ tumors, using two independent methods of predicting macrophage polarization ^32,33^, macrophages within pnSTING^low^ tumors were predicted to be “M2” -like or alternatively activated, suggesting an infiltrating pro-tumorigenic myeloid population (**Fig. 3c**). This is consistent with reports that activation of STING signaling improves activated: inhibitory ratios of tumor-associated macrophages, resulting in an anti-tumor immune response ^34^.

**Figure 4:**
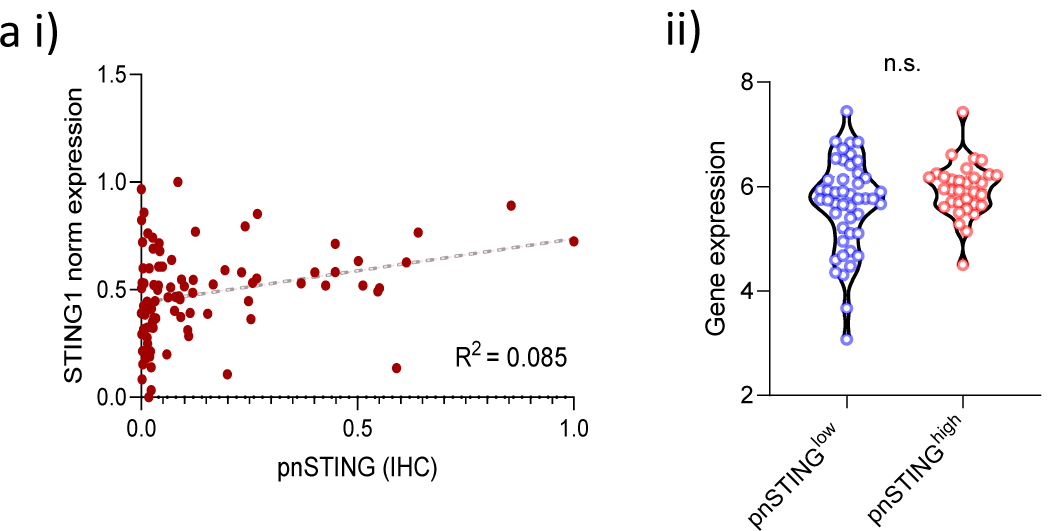
pnSTING and STING1 gene expression. **a i**) Correlation plot of normalized pnSTING score (IHC) and STING1 gene expression. R = 0.085. **ii**) Gene expression of STING1 compared between pnSTING^low^ (< median) and pnSTING^high^ (> median) ER+ breast cancers. n.s. = non significant.

### Identification of a gene signature characterizing pnSTING^low^ ER+ poor prognosis tumors

The poor outcomes in pnSTING^low^ ER+ tumors, a typically good prognosis subtype of breast cancer, led us to explore the molecular characteristics defining this subgroup. To characterize pnSTING^high^ and pnSTING^low^ ER+ tumors, we performed GSEA which identified a subset of genes enriched in both (**Suppl. Table 4**). As the genes associated with pnSTING^high^ were predominately immune related and likely derived from the immune-infiltrated TME, we chose to focus on the genes upregulated in pnSTING^low^. Using the top 25 genes altered in pnSTING^low^ tumors, stratification based on high or low gene signature (> median <), replicated the survival differences observed when tumors were stratified based high and low perinuclear STING IHC expression (**Suppl. Fig. 4a-b**). To validate this further, we interrogated 4 independent datasets using the pnSTING^low^ signature. These included: independent samples from the discovery cohort that were not available for IHC-based STING analysis with mRNA expression data available, METABRIC, TCGA 2012 and Wang ^22,23,26^

Consistently in each independent dataset we found the pnSTING^low^ signature predicted poor survival in ER+ cases (**Fig. 5a-d, Suppl. Table 5**). Exploring clinicopathological and genomic correlates within the METABRIC and TCGA datasets, ER+ pnSTING^low^-signature cases were associated with increased tumor grade (**Fig. 5e**) as well as Luminal B-like cases (**Fig. 5fi**). Increased chromosomal instability, as measured by copy number gain and fraction genome altered, was associated with pnSTING^low^-signature cases (**Fig. 5fii, Suppl. Fig. 5eii**). Other significant variations are reported in **Suppl. Fig. 5**. Importantly, although all samples in the discovery dataset received chemotherapy treatment, suggesting a higher risk population, within the METABRIC cohort we identified little variation in rates of chemotherapy in pnSTING^low^-signature cases (**Fig. 5eiii**). This was confirmed in the Wang dataset, where all patients received hormone therapy without chemotherapy. Therefore, we have identified a signature of poor prognosis in ER+ breast cancer independent of systemic anti-cancer therapy received, suggesting alternative approaches should be considered in this subgroup.

**Figure 5:**
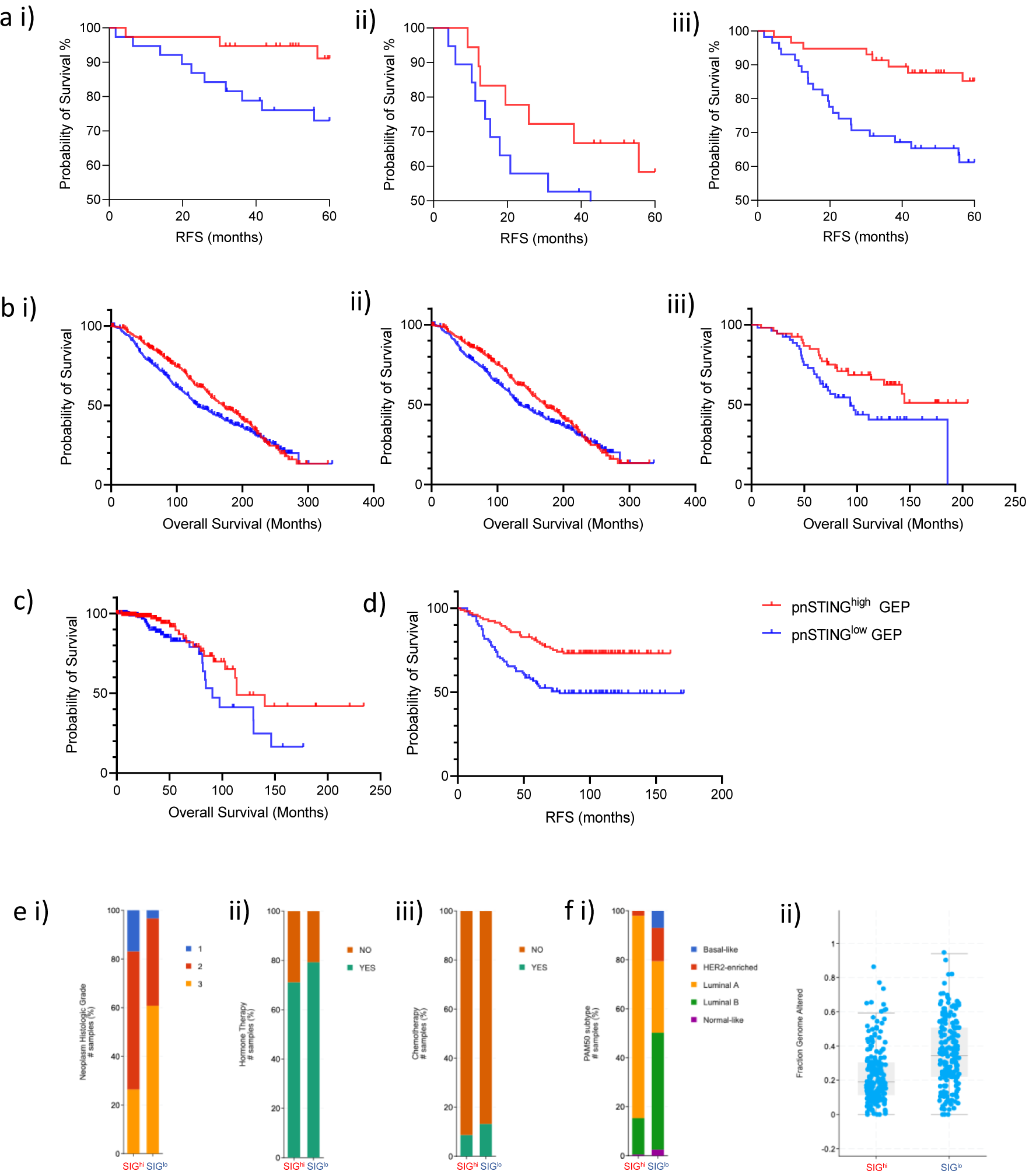
pnSTING^low^ gene expression signature is prognostic of poor outcome in independent datasets. **a**) Kaplan Meier Curve analysis of relapse free survival (RFS - months) in **i**) ER positive (ER+), **ii**) ER negative (ER-) and **iii**) all samples with gene expression data only from discovery dataset stratified by pnSTING signature score. **b**) Kaplan Meier Curve analysis of overall survival (months) in **i**) ER+ disease (all treatments) **ii**) those receiving hormone therapy only **iii**) those receiving chemotherapy in the METABRIC dataset stratified by pnSTING signature score. **c** & **d**) Kaplan Meier Curve analysis of overall survival (months) in ER+ disease from the **c**) TCGA 2012 dataset and **d**) Wang dataset stratified by pnSTING signature score. **e**) Clinicopathologial and molecular characteristics of METABRIC samples classified by pnSTING gene expression score comparing **i**) Tumor grade (adj. p <0.0001) **ii**) Hormone therapy (adj. p = 0.009) **iii**) Chemotherapy (adj. p = 0.048) **f**) Clinicopathologial and molecular characteristics of TCGA samples classified by pnSTING gene expression score comparing **i**) PAM50 subtype (adj. p <0.0001) and **ii**) Fraction genome altered (adj. p < 0.0001).

### Targetable pathways within pnSTING^low^ poor prognosis tumors

Given the consistently poor outcomes observed in ER+ pnSTING^low^ breast cancer, we sought to identify potentially targetable pathways. To do so, we utilized multi-omic data in the discovery and validation cohorts. We noted that ER+ pnSTING^low^ tumors were associated with genesets associated with chromosomal instability (**Fig. 6a-b**). Further examination of the individual genes within the signature revealed upregulation of chromatin regulation and DNA repair genes *CDT1, NCAPD3* and *EXO1*. The top gene in this signature, *IMPA2*, has recently been reported to drive cervical cancer progression with a potential novel role in DNA repair ^35^.

**Figure 6:**
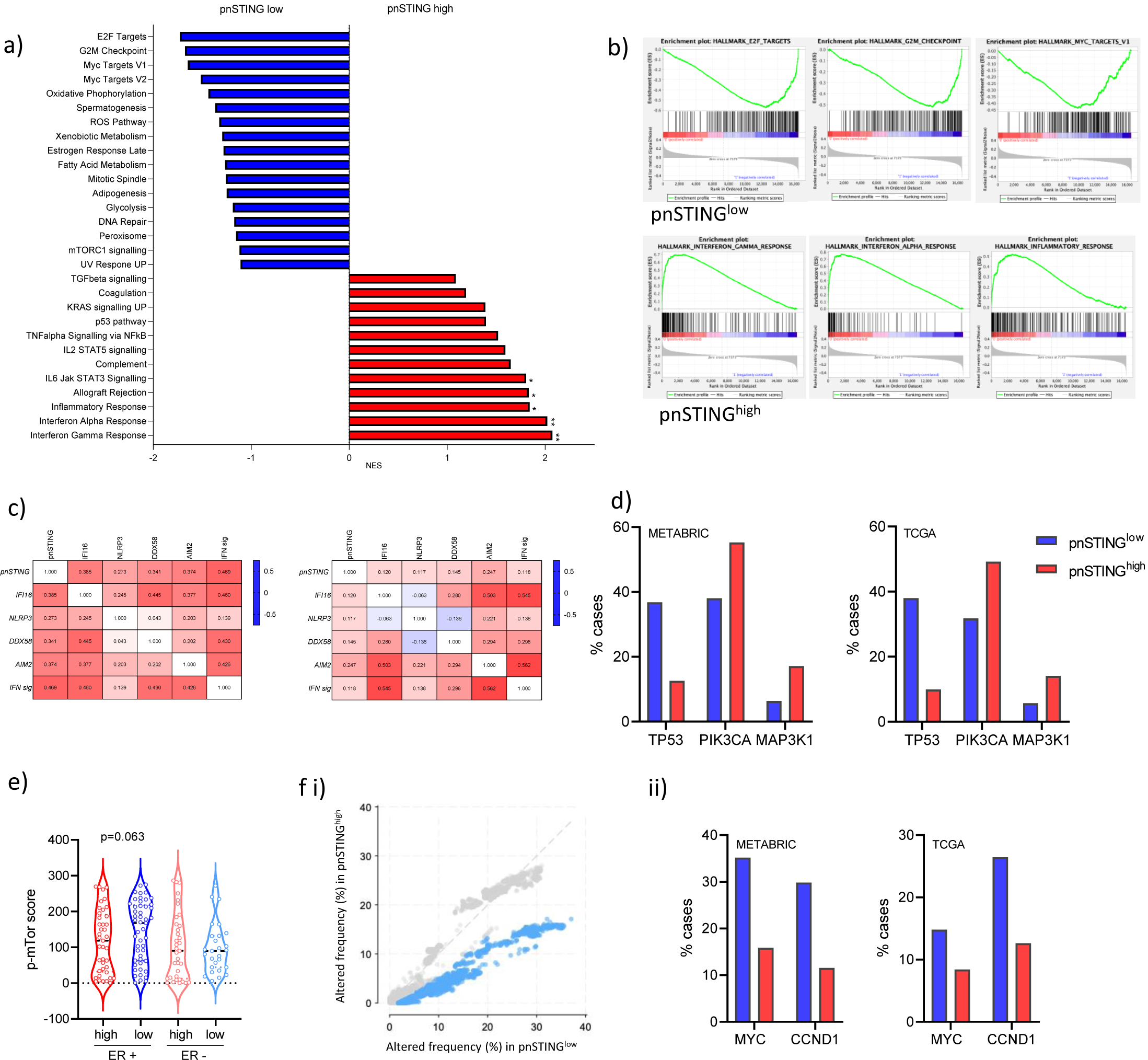
Genomic and transcriptomic analysis of pnSTING signature classified ER+ tumors. **a**) GSEA results showing Normalised Enrichment Scores (NES) for the gensets enriched > ± 1 in pnSTING^high^ signature (red) or pnSTING^low^ signature (blue) samples. **b**) Enrichment plots for the top 3 genesets enriched in pnSTING^high^ signature or pnSTING^low^ signature samples. **c**) Correlation matrices for (left) ER positive and (right) ER negative breast cancer for pnSTING value, gene expression of DNA and RNA sensors and 18-gene interferon signature. **d**) Percentage of ER+ cases with mutations in *TP53, PIK3CA* and *MAP3K1* in the METABRIC (left) and TCGA (right) 2012 datasets stratified based on high (above median) and low (below median) STING signature score. **e**) Expression of p-mTOR (Ser2448) quantified using QuPath in the discovery dataset stratified based on ER expression and perinuclear STING expression (where red = pnSTING^high^ and blue = pnSTING^low^). **f**) Percentage of ER+ cases with copy number alteration **i**) overall (with significance indicated in blue) in the METABRIC dataset and **ii**) *MYC* and *CCND1* in the METABRIC and TCGA 2012 datasets stratified based on high (above median) and low (below median) STING signature score.

In contrast to ER+ breast cancer, there was an unexpected disconnection between pnSTING and immune cell infiltration in ER-disease, as well as reversal in the prognostic ability of the pnSTING^low^ signature in independent ER-datasets (**Suppl. Fig. 4c-e**). We analyzed the correlation between pnSTING score (based on IHC data) and gene expression of other cytosolic DNA sensors, as well as an 18-gene T-cell inflamed signature identifying active interferon responses^36^ (**Fig. 6c, Suppl. Fig. 6ai-ii**). Importantly, while we observed an expected correlation between STING, other nucleic acid sensors and interferon signaling in ER+ disease, this was notably absent in ER-disease, with no significant correlation observed between pnSTING and interferon activity. As STING has important non-interferon related functions^37,38^, we hypothesize that post-translational modifications of STING may result in immune-independent functions being the dominant role of STING in ER-breast cancer.

Using the available somatic mutation and copy number alteration data associated with repressed STING signaling in the ER+ METABRIC and TCGA 2012 cohorts, we identified high rates of *TP53* mutations (36.8% vs 12.6%; 38.0% vs 9.95% respectively) (**Fig. 6d, Suppl. Table 6**). Interestingly, a recent report identifies suppression of STING signaling via mutant p53 binding and inhibition of TBK1 downstream of STING in breast cancer cells, suggesting that STING-mediated interferon responses may be prevented in p53 mutant cancer cells via this mechanism ^39^. In contrast, higher rates of mutation in *PI3K* and *MAPK1* were observed in pnSTING^high^ signature cases. *PI3K* mutations have been reported to correlate with Luminal A cancers, and good prognosis, consistent with our data ^40^. In the same report, mTOR pathway activation was associated with Luminal B subtype and poor outcomes. Given that GSEA also identified increased mTOR signaling in pnSTING^low^ samples, we sought to validate this finding by IHC analysis of phosphorylated mTOR in the discovery dataset (**Suppl. Fig. 6b**). Although not significant, a trend towards increased levels of p-mTOR in the pnSTING^low^ cohort was noted, consistent with GSEA (p = 0.063, **Fig. 6e**). mTOR inhibitors have previously been studied in ER+ breast cancer, but have lacked a specific biomarker predicting response, and further study of this pathway in pnSTING^low^ tumors may indicate a means of selecting patients for this therapy.

Rates of copy number alteration were significantly increased in pnSTING^low^-signature ER+ cases in the METABRIC dataset, although this did not reach significance in the TCGA dataset (**Fig. 6fi, Suppl. Fig. 6c**). In particular, regions encoding *MYC* (8q24) and *CCND1* (11q13) were significantly amplified in pnSTING^low^ cases in both cohorts (**Fig. 6fii, Suppl. Table 7**). *CCDN1* amplification, encoding CyclinD1, results in chromosomal instability via *CDT1* ^41^, consistent with our gene expression findings, suggesting that CDK4/6 inhibitors may also have a role for the treatment of this subgroup of breast cancer. *MYC* amplification was consistent with GSEA identification of upregulation of MYC targets in pnSTING^low^ tumors. Via the ENCODE database, we identified predicted binding sites for MYC and the co-factor MAX (Myc-associated factor X) within *STING* (**Suppl. Fig. 7**), although it is not known whether MYC directly regulates STING or indirectly regulates downstream responses.

## Discussion

Understanding the regulation of STING-induced immune responses is crucial to understanding immune evasion and improving the anti-cancer effectiveness of immune-targeting and other therapies. By developing a novel digital pathology approach assessing STING on IHC sections, and subsequently identifying a gene signature to classify pnSTING^high/low^ cancers, we were able to validate our finding of poor prognosis in pnSTING^low^ ER+ breast cancer in both chemotherapy and hormone therapy-treated tumors. Interestingly, while there was an observed connection between pnSTING and immune infiltration in ER+ breast cancers, this was not the case in ER-disease. Previous reports have identified the role of tumor infiltrating lymphocytes in predicting responses to chemotherapy in triple negative and HER2 positive breast cancers, but the role in ER+ disease has been less well-defined ^42,43^. This suggests that measuring pnSTING may be able to add granularity to the activation state of the immune infiltrate in ER+ disease, therefore identifying tumors with immune restriction and poor clinical outcomes to standard of care, suggesting alternative treatment options should be considered.

Importantly we identified *MYC* amplification associated with chromsomal instability as a potential mechanism of STING repression. Typically, *MYC* is amplified in 15% of breast cancers, and associated with resistance to hormone therapy in Luminal A cancers ^44^. We found *MYC* amplification in up to 35% of pnSTING^low^ ER+ breast cancers, a significant enrichment in this subgroup. Amplification of *MYC* is commonly associated with, and a driver of, chromosomal instability ^45^. As chromosomal instability results in cGAS stimulation via micronuclei and the subsequent production of 2’3’cGAMP, it is logical that cancer cells will develop mechanisms to repress anti-cancer immune responses induced by STING. Interestingly MYC has recently been reported to suppress STING signaling in *TP53*- and *BRCA1*-mutant breast cancer ^46^. This supports the hypothesis that amplification of *MYC* is a mechanism of STING repression (and subsequent immunosuppression) in pnSTING^low^ tumors, potentially targetable using novel MYC inhibitors. Indeed, MYC complexes bind directly to IRF5, IRF7, STAT1 and STAT2 in pancreatic cancer, repressing interferon responses ^47^. MYC may have a dual role in cancer progression, promoting both chromosomal instability and direct or indirect suppression of STING-mediated immune responses, resulting in an immunosuppressed microenvironment. The use of novel MYC inhibitors has been shown to increase immune infiltration in preclinical models although the mechanisms driving this infiltrate have not yet been described, and may be STING related ^48^. Combining MYC inhibition and STING activation requires further preclinical study and could be a therapeutic approach in pnSTING^low^ cases.

Other genomic alterations identified in pnSTING^low^ ER+ cases include *CCND1* amplification. There are early reports linking CCND1 amplification to immunosuppressive signaling and resistance to ICB ^49^. Interestingly CDK4/6 inhibitors synergize with ICB to enhance responses ^50^, an approach which is now being tested in early phase clinical trials. Additionally, the finding of increased mTOR activity in the pnSTING^low^ subgroup may suggest an alternative therapeutic approach. mTOR has complex and varying roles in the TME, including “M2” macrophage polarization and promoting myeloid derived suppressor cell accumulation via upregulation of G-CSF ^51^. In keeping with this, mTOR inhibition demonstrates increased activation and persistence of intratumoral T-cells ^52^.

Our study focused on early stage breast cancer, and further work is needed to determine if this chromosomally unstable pnSTING^low^ subgroup persists through metastatic disease. Moreover, we did not identify a clear link between pnSTING expression and outcomes in ER-disease. ER-breast cancer more commonly involves mutation of DNA repair genes including *BRCA1/2, PALB2* and members of the Fanconi Anemia pathway, which result in STING pathway activation. While these DNA repair defects result in aggressive disease, they are vulnerable to DNA-damaging chemotherapy which is standard in the treatment of triple negative breast cancer. This paradox may explain the conflicting findings in ER+ and ER-breast cancer, although further study is needed. In contrast to ER+ disease, we identified an uncoupling of STING expression and interferon responses in ER-breast cancer. The reasons for this are currently unclear – it may be that STING-modulating mechanisms such as post-translational modifications drive STING towards non-interferon signaling that promotes cancer progression.

While we observed STING staining in the stroma of tumors, we were unable to delineate expression in fibroblasts v. immune cells, for example. Recently, specific efflux and/or influx channels for 2’3’cGAMP have been identified for monocytes, macrophages, fibroblasts and endothelial cells ^53–57^. As STING activation within host immune cells is key for anti-cancer responses, expression of these channels within cancers may drastically alter the nature of STING responses in response to either endogenous or exogeneous STING activation. Expression of STING within the stroma may reflect presence or absence of these transport channels, and further modulate immune responses. Whether these channels differ between ER+ and ER-cancers is currently not known. Moreover, cGAS-STING pathway activation within fibroblasts can promote resistance to oncolytic viral therapy, with sensitivity restored by TBK1 inhibition, suggesting STING signaling within fibroblasts may indeed promote tumor progression ^58^. Further exploration of this may address the unexpecting finding of a disconnection between STING and interferon responses in ER-disease.

In summary, taking a novel digital pathology IHC-based approach, we can identify tumors with high intrinsic STING activation. Taken together with identification of immune infiltration, this may identify tumors which respond favorably to immune checkpoint blockade and/or STING agonists. Further exploration of this assay in the context of immune modulating treatment and metastatic disease is warranted. By utilizing multiomic data from independent breast datasets, we have characterized pnSTING^low^ ER+ cancers as a subgroup of breast cancer with intrinsic immunosuppression and chromosomal instability, with poor response rates to standard chemotherapy or endocrine therapy. Further study of pathways resulting in immunosuppression and potential alternative approaches in this subgroup may result in improved patient outcomes.

## Acknowledgments

The samples used in this research were received from the Northern Ireland Biobank which has received funds from HSC Research and Development Division of the Public Health Agency in Northern Ireland and the Friends of the Cancer Centre. The Northern Ireland Molecular Pathology Laboratory was responsible for construction of tissue microarrays, slide staining, and scanning.

EEP was supported by the Academy of Medical Sciences (Starter Grant for Clinical Lecturers), the Prostate Cancer Foundation (Young Investigator Award) and the Oxford Institute for Radiation Oncology.

MPH, SGC and MST were supported by CRUK Accelerator (C11512/A20256). We are grateful to the NVIDIA Corporation for supporting our research via the GPU Grant Program for researchers.

SMP was supported by the British Research Council.

NEB was supported by Breast Cancer Now Scientific Fellowship (2012MaySF122).

## Author contributions

Conception: EEP, NEB, MPH, MST, SMcQ. Data acquisition: EEP, MPH, EG, FAS, VB, SGC, SMP, CG, JM, DG, SMcQ, MST, NEG. Data analysis: EEP, MPH, EG, RDK, SFB, SMcQ, MST, NEB.

Manuscript writing and editing: EEP, MPH, RDK, SFB, MST, NEB. Confirmation of final draft: All authors.

