## Supplementary Figures for "The clinical and molecular significance associated with STING signaling in estrogen receptor-positive early breast cancer"

*Suppl. Figure 1: Tumor, Stromal and Perinuclear STING expression in breast cancer*

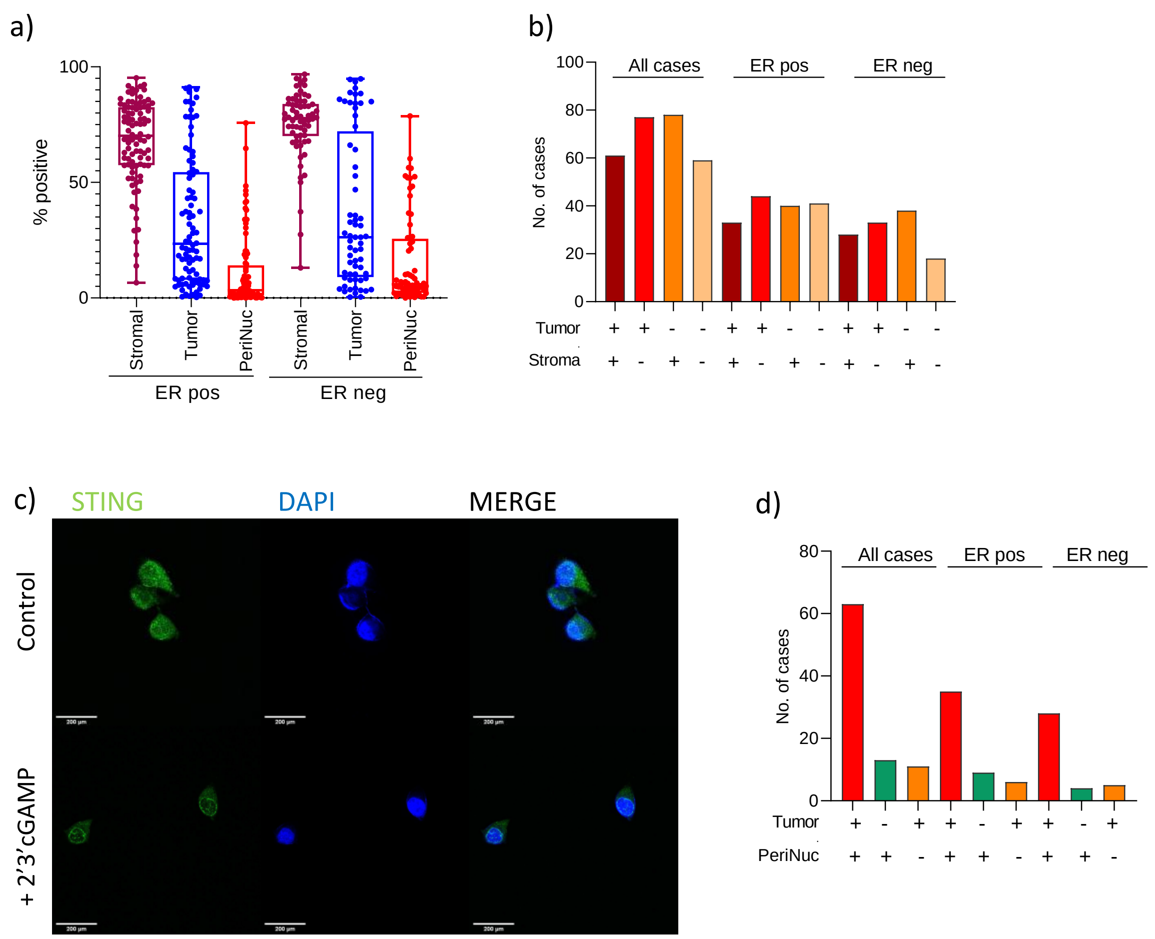


*Supplementary Figure 1* **a**) Percentage positive cells for stromal, tumor and perinuclear STING in ER positive and ER negative tumors in discovery dataset. **b**) Distribution of tumor and stromal expression of STING where + represents > median expression and - represents < median expression. Median is calculated on whole dataset expression. **c**) Immunofluorescence images of MDA436-EV cells treated with either control or 2'3' cGAMP (10 µg/ml) for 60 minutes. Scale bar = 200 µm. **d**) Unique expression of tumor or perinuclear-only STING in all cases, ER positive and ER negative breast tumors where + represents > median expression and - represents < median expression. Median is calculated on whole dataset expression.

*Suppl. Figure 2: pnSTING IHC score predicts outcome in ER+ breast cancer*

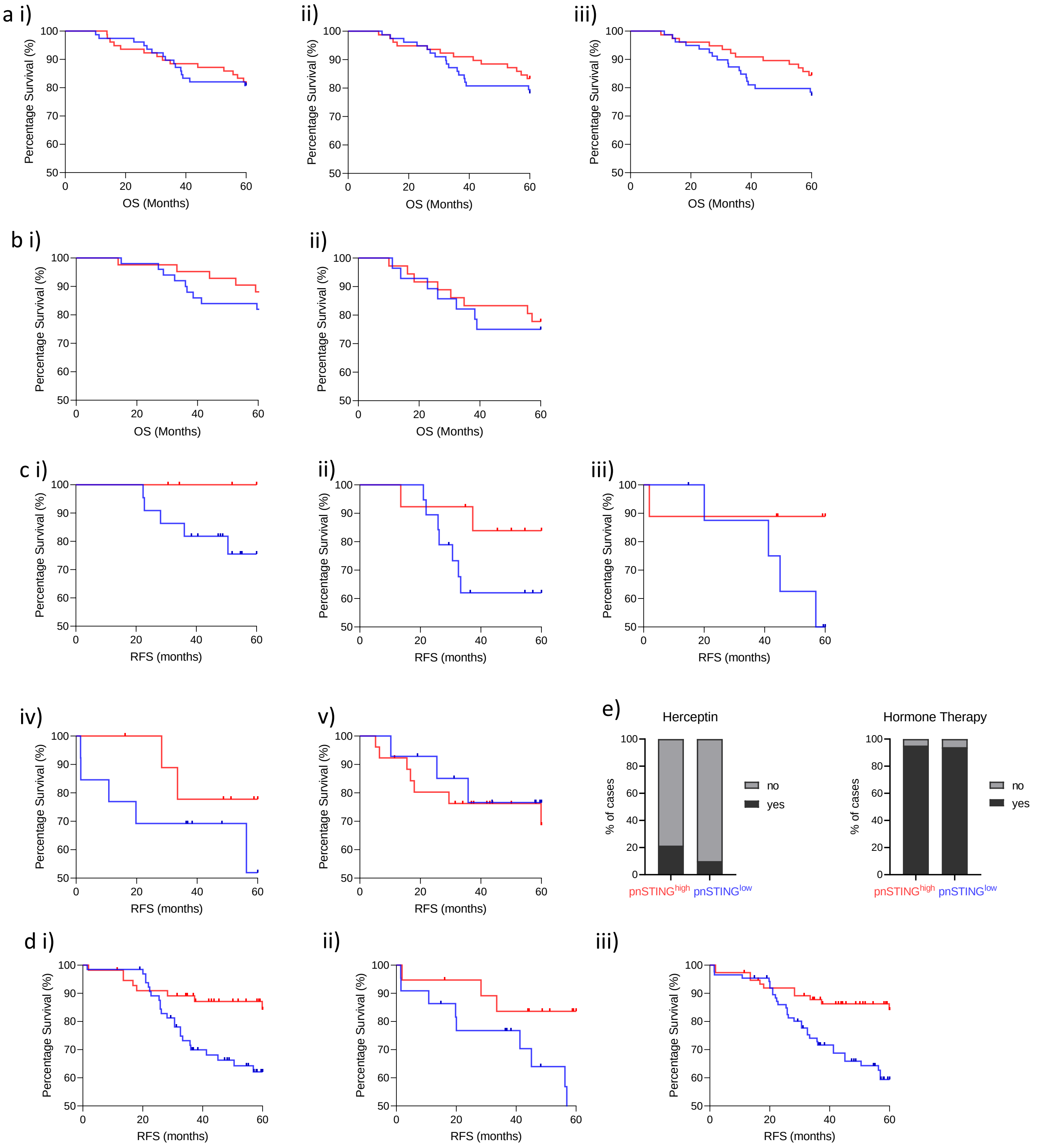


*Supplementary Figure 2* **a**) Kaplan Meier Curve of overall survival (OS) stratified based on high (above median) or low (below median) of STING expression in the **i**) stromal compartment, **ii**) tumour epithelial compartment or **iii**) perinuclear region. **b**) Kaplan Meier Curve of overall survival (OS) stratified based on high (above median of all cases) or low (below median of all cases) of STING expression in the perinuclear region in **i**) ER positive (ER+) and **ii**) ER negative (ER-) cases. **c**) Kaplan Meier Curve of Relapse Free Survival (RFS) stratified based on high (above median) or low (below median) of STING expression in the perinuclear region in St Gallen subtypes: **i**) ER+/HER2-/Ki67-, **ii**) ER+/HER2-/Ki67+, **iii**) ER+/HER2+/Ki67+, **iv**) ER-/HER2+ and **v**) ER-/PR-/HER2- (TNBC). **d**) Kaplan Meier Curve of Relapse Free Survival (RFS) stratified based on high (above median) or low (below median) of STING expression in the perinuclear region in **i**) ER+/HER2-/Ki67+/- , **ii**) ER+/-/HER2+and **iii**) Non TNBC. **e**) Stacked bar chart of the percentage of ER+ patients stratified based on high (above median) or low (below median) of STING expression in the perinuclear region based on clinical attributes: Herceptin Treatment (left panel), Hormone Therapy (right panel).

*Supplementary Figure 3: pnSTING immune correlates in ER+ and ER- breast cancer*

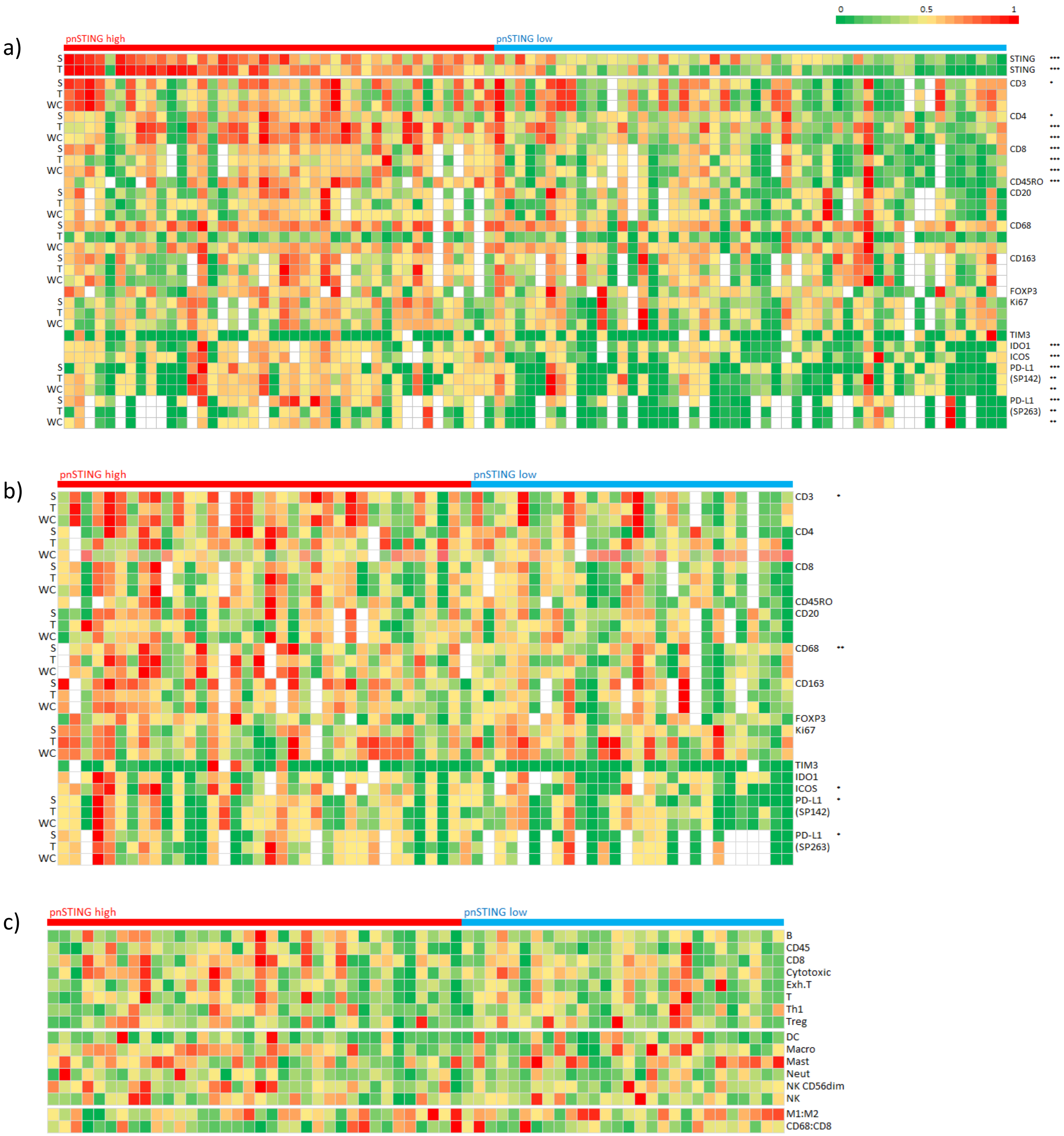


*Supplementary Figure 3* **a**) Heatmap of normalized expression measured by IHC of perinuclear STING, stroma STING, tumor STING, CD3, CD4, CD68, CD45RO, CD163, FOXP3, CD8, CD20, Ki-67, PD-L1 measured by SP263 and SP142, TIM3, IDO1 and ICOS in ER+ breast cancer cases. Expression was quantified in the stroma compartment (S), tumor epithelial compartment (T) or whole core (WC) as indicated. **b**) Heatmap of normalized expression measured by IHC of perinuclear STING, stroma STING, tumor STING, CD3, CD4, CD68, CD45RO, CD163, FOXP3, CD8, CD20, Ki-67, PD-L1 measured by SP263 and SP142, TIM3, IDO1 and ICOS in ER- breast cancer cases. Expression was quantified in the stroma compartment (S), tumor epithelial compartment (T) or whole core (WC) as indicated. **c**) Heatmap of normalized immune scores derived from deconvolution of microarray data in ER- breast cancer cases. Correlation between markers and perinuclear STING stratified based on high (above median) and low (below median) using the Krushall Wallis test on non-transformed data with *, ** and *** indicating p values of $< 0.05$, $< 0.01$ and $< 0.001$ respectively.

*Suppl .Figure 4: pnSTING$^{low}$ gene expression signature in discovery dataset and ER negative breast cancer*

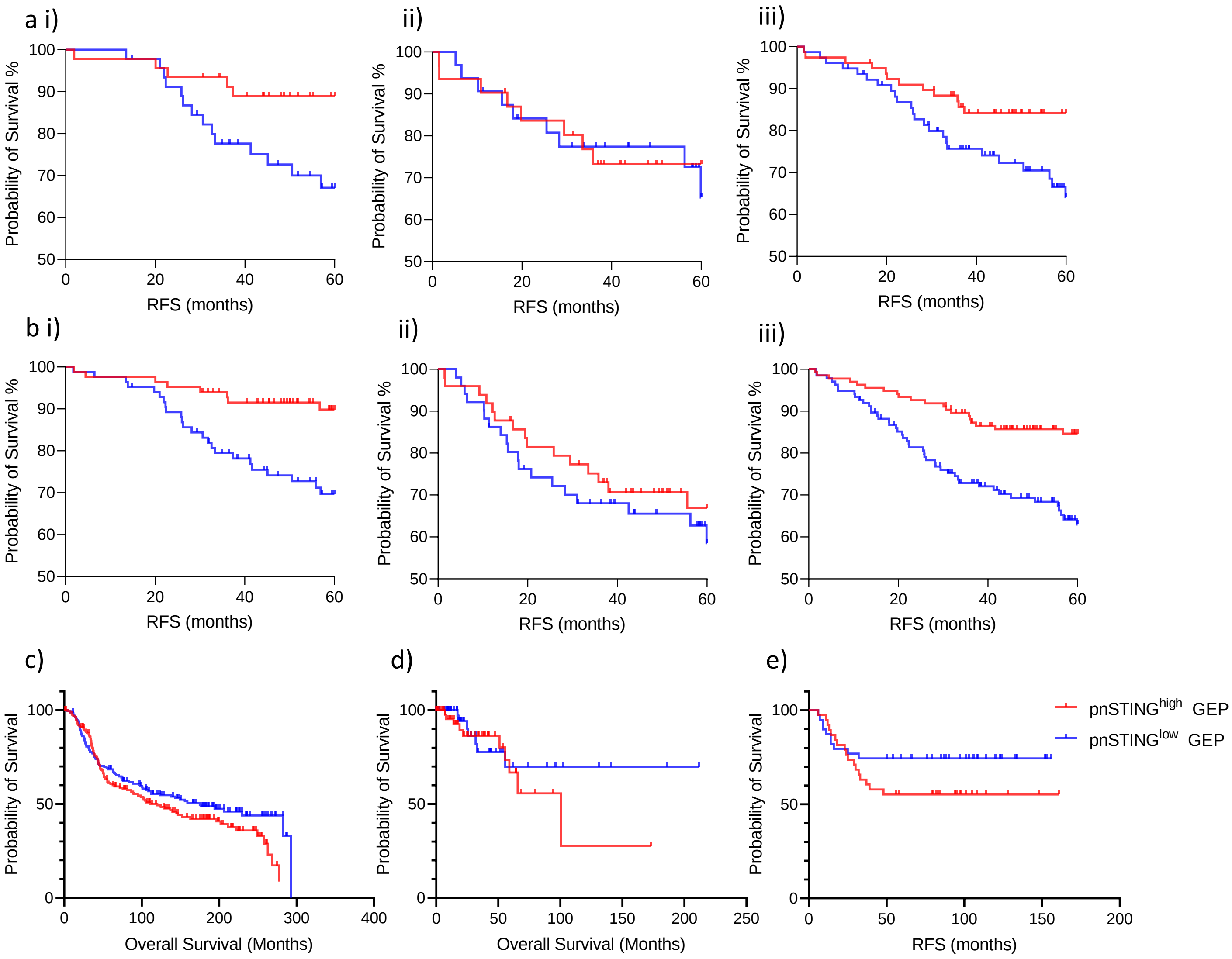


*Supplementary Figure 4* **a**) Kaplan Meier Curve analysis of relapse free survival (RFS - months) in **i**) ER positive (ER+), **ii**) ER negative (ER-) and **iii**) all samples with STING IHC data from discovery dataset stratified by pnSTING signature score. **b**) Kaplan Meier Curve analysis of relapse free survival (RFS - months) in **i**) ER positive (ER+), **ii**) ER negative (ER-) and **iii**) all samples with STING IHC and/or gene expression data from discovery dataset stratified by pnSTING signature score. **c-e**) Kaplan Meier Curve analysis of overall survival (months) in ER- disease from the **c**) METABRIC **d**) TCGA 2012 and **e**) Wang datasets.

*Suppl .Figure 5: Clincopathological and molecular characteristic of pnSTING$^{high}$ and $^{-low}$ breast cancers*

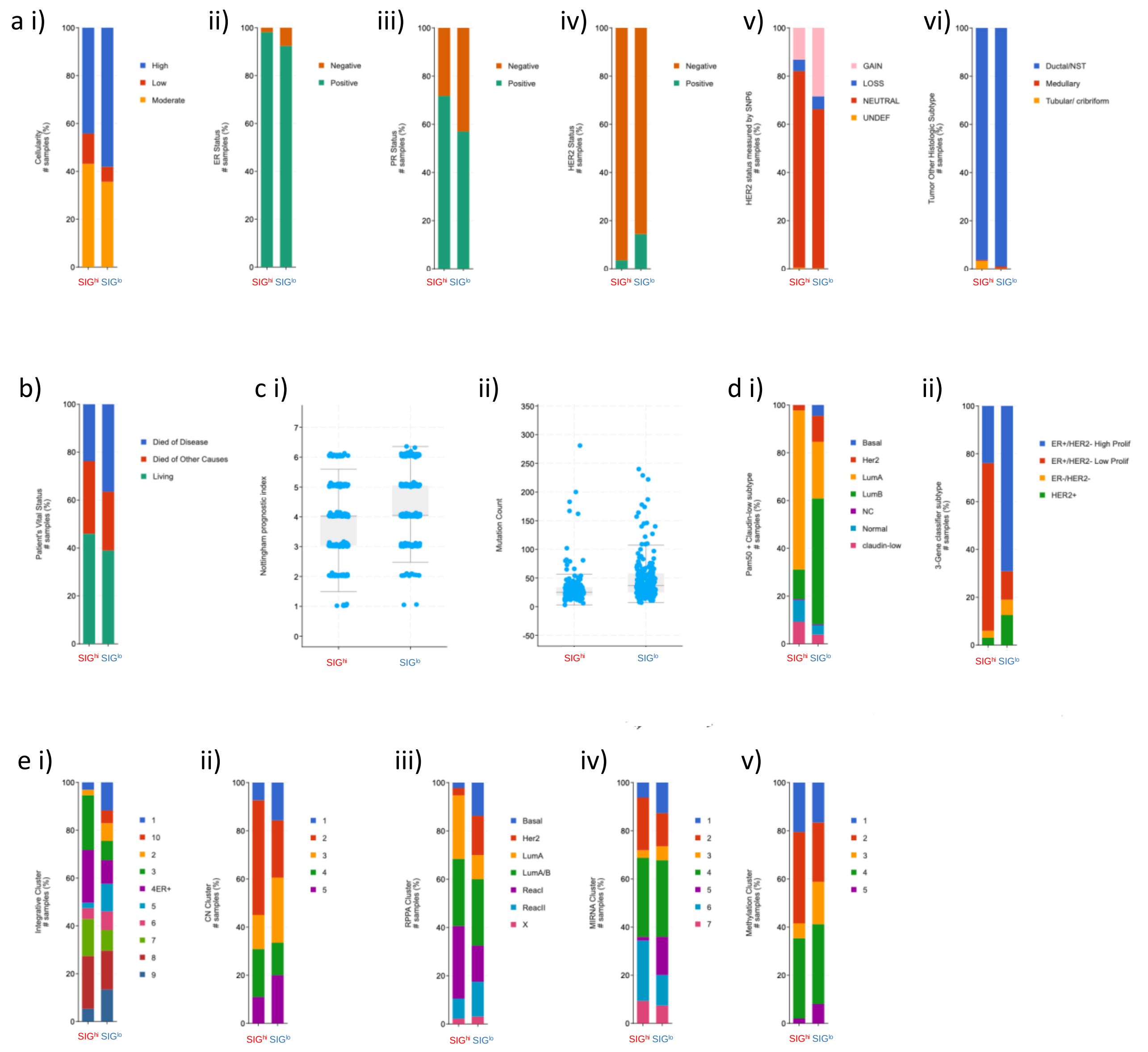


*Supplementary Figure 5* METABRIC and TCGA ER+ cohort data stratified by pnSTING signature score for **a i**) Cellularity **ii**) ER status **iii**) PR status **iv**) HER2 status **v**) HER2 status by SNP6 measurement **vi**) histological subtype. **b**) Patient outcomes. **c i**) Nottingham prognostic index score (all METABRIC) **ii**) Mutational load (TCGA) **d i**) PAM50 and Claudin low classified subgroups **ii**) 3-gene classifier (METABRIC) **e i**) Integrative Clusters **ii**) CN clusters **iii**) RPPA **iv**) miRNA clusters and **v**) Methylation clusters (TCGA).

*Suppl. Figure 6: mTOR expression and pnSTING correlations*

a i)

| | pnSTING | IFI16 | NLRP3 | DDX58 | AIM2 | IFN sig |
|---|---|---|---|---|---|---|
| **pnSTING** | | 1.51E-04 | 8.35E-03 | 8.84E-04 | 2.41E-04 | 2.4E-06 |
| **IFI16** | 1.51E-04 | | 1.86E-02 | 9.00E-06 | 2.08E-04 | 3.9E-06 |
| **NLRP3** | 8.35E-03 | 0.0186 | | 0.685 | 0.0521 | 0.186 |
| **DDX58** | 8.84E-04 | 9.00E-06 | 0.685 | | 0.0534 | 1.9E-05 |
| **AIM2** | 2.41E-04 | 2.08E-04 | 0.052 | 0.0534 | | 2.3E-05 |
| **IFN sig** | 2.36E-06 | 3.89E-06 | 0.186 | 1.90E-05 | 2.25E-05 | |

ii)

| | pnSTING | IFI16 | NLRP3 | DDX58 | AIM2 | IFN sig |
|---|---|---|---|---|---|---|
| **pnSTING** | | 0.344 | 0.357 | 0.254 | 0.0493 | 0.354 |
| **IFI16** | 0.344 | | 0.619 | 0.025 | 2.25E-05 | 3.24E-06 |
| **NLRP3** | 0.357 | 0.619 | | 0.285 | 0.0787 | 0.278 |
| **DDX58** | 0.254 | 0.025 | 0.285 | | 0.0184 | 0.017 |
| **AIM2** | 0.049 | 2.25E-05 | 0.079 | 0.018 | | 1.38E-06 |
| **IFN sig** | 0.354 | 3.24E-06 | 0.278 | 0.017 | 1.38E-06 | |

b i) ii) iii) iv)

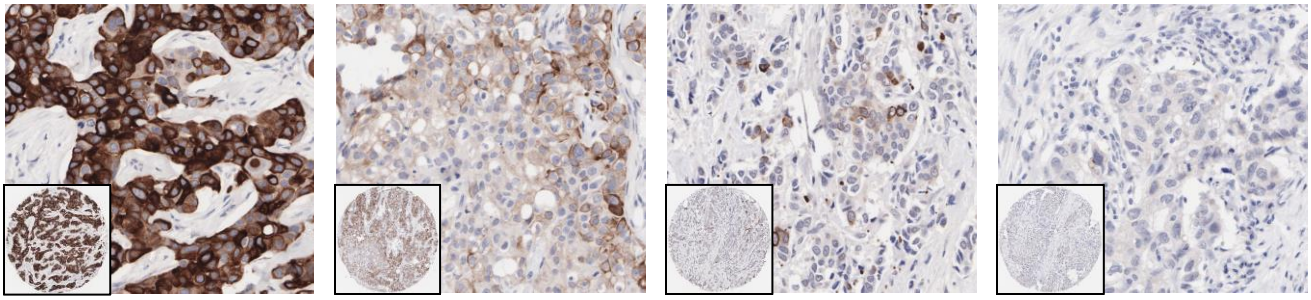

c)

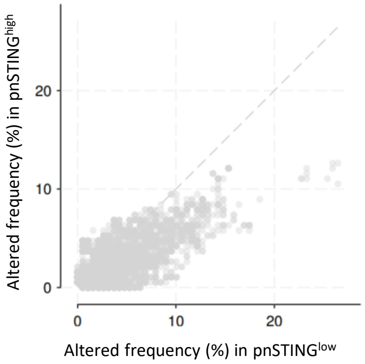


*Supplementary Figure 6* **a**) P values calculated correlations in **i**) ER positive (left table) and **ii**) ER negative (right table) breast cancers for pnSTING values, gene expression of cytosolic DNA and RNA sensors and interferon signaling. Non-significant values are indicated in blue. **b**) Representative IHC images of breast cancer samples with **i**) high, **ii**) medium, **iii**) low and **iv**) absent mTOR signaling using phospho-specific mTOR antibody (Ser2448). Magnification x 40, Inset x 10. **c**) Percentage of ER+ cases with copy number alteration overall in the TCGA dataset.

*Suppl. Figure 7: Transcription factor analysis of STING1 using ENCODE database*

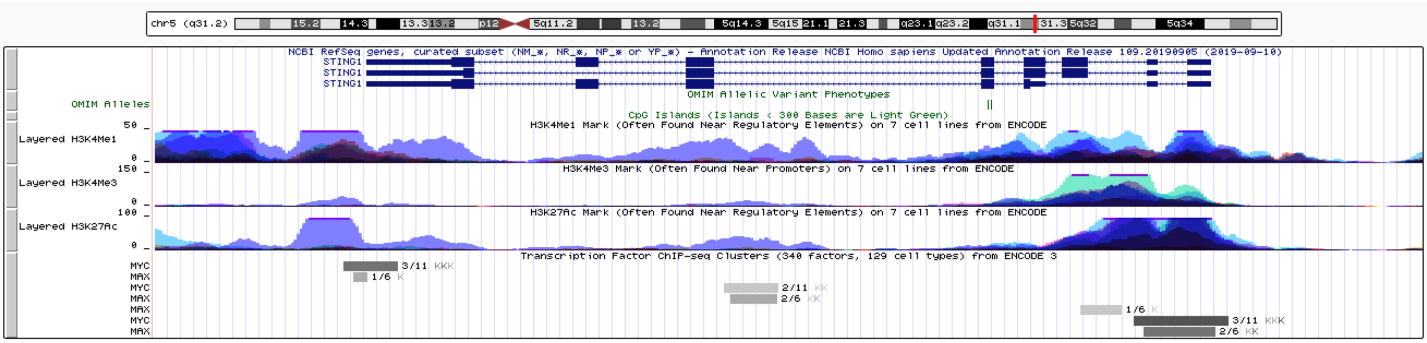


*Supplementary Figure 7* MYC and MAX predicted binding sites on *STING1*. (Note: *STING1* is encoded on the negative strand and therefore is viewed from right to left.)
