## Supplementary Tables for "The clinical and molecular significance associated with STING signaling in estrogen receptor-positive early breast cancer"

*Supplementary Table 1:* Antibodies used in IHC

| *Antibody* | *Clone* | *Company* | *Catalogue Number* | *Platform* | *Retrieval* | *Dilution* | *Incubation* | *Detection* |
|---|---|---|---|---|---|---|---|---|
| ***STING*** | Poly | Protein Tech | 19851-1-AP | Leica Bond RX | ER1 20min<br>ER1 20min | 1:1000<br>1:2000 | 15min @ RT<br>30min @ RT | DAB Polymer Refine + Enhancer<br>Opal Fluorophore 690 |
| ***ER*** | 6F11 | Leica | NCL-L-ER-6F11 | Leica Bond RX | ER2 30min | 1:200 | 15min @ RT | DAB Polymer Refine + Enhancer |
| ***HER2*** | CB11 | Leica | NCL-L-CB11 | Leica Bond RX | ER1 25min | 1:40 | 15min @ RT | DAB Polymer Refine + Enhancer |
| ***KI-67*** | 30-9 | Roche | 790-4286 | Ventana Benchmark XT | CC1 32min | Neat | 16min @ 37C | Optiview DAB Kit |
| ***CK*** | AE1/AE3 | Dako | M3515 | Leica Bond RX | ER2 20min<br>ER2 20min | 1:200<br>1:200 | 15min @RT<br>30min @RT | DAB Polymer Refine + Enhancer<br>Opal Fluorophore 520 |
| ***CD3*** | 2GV6 | Roche | 790-4341 | Ventana Benchmark XT | CC1 32min | Neat | 16min @ 37C | Optiview DAB Kit |
| ***CD4*** | SP35 | Roche | 790-4423 | Ventana Benchmark XT | CC1 32min | Neat | 16min @ 37C | Ultraview DAB Kit |
| ***CD8*** | C8/144B | Dako | M7103 | Leica Bond RX | ER2 20min | 1:200 | 15min @ RT | DAB Polymer Refine + Enhancer |
| ***CD20*** | L26 | Dako | M0755 | Leica Bond RX | ER1 30min | 1:400 | 15min @ RT | DAB Polymer Refine + Enhancer |
| ***CD68*** | 514H12 | Leica | PA0273 | Leica Bond RX | ER2 20min | 1:200 | 30min @ RT | DAB Polymer Refine + Enhancer |
| ***CD45RO*** | UCHL1 | Leica | NCL-L-UCHL1 | Leica Bond RX | ER1 20min | 1:2000 | 15min @ RT | DAB Polymer Refine + Enhancer |
| ***CD163*** | 10D6 | Leica | PA0090 | Leica Bond RX | ER2 20min | Neat | 15min @ RT | DAB Polymer Refine + Enhancer |
| ***FOXP3*** | SP97 | LSBIO | LS-C210349 | Leica Bond RX | ER2 20min | 1:50 | 15min @ RT | DAB Polymer Refine + Enhancer |
| ***ICOS*** | D1K2T | Cell Signaling | #89601 | Leica Bond RX | ER2 20min | 1:400 | 15min @ RT | DAB Polymer Refine + Enhancer |
| ***IDO1*** | D5J4E | Cell Signaling | #86630 | Leica Bond RX | ER2 20min | 1:400 | 15min @ RT | DAB Polymer Refine + Enhancer |
| ***mTOR*** | 49F9 | Cell Signaling | #2976 | Leica Bond RX | ER2 20min | 1:100 | 15min @ RT | DAB Polymer Refine + Enhancer |
| ***PD-L1*** | SP263 | Roche | 790-4905 | Ventana Benchmark XT | CC1 64min | Neat | 16min @ 37C | Optiview DAB Kit |
| ***PD-L1*** | SP142 | Spring Bioscience | M4420 | Ventana Benchmark XT | CC1 64min | 1:40 | 16min @ 37C | Optiview DAB Kit |
| ***TIM3*** | D5D5R | Cell Signaling | #45208 | Leica Bond RX | ER2 20min | 1:200 | 15min @ RT | DAB Polymer Refine + Enhancer |

*Supplementary Table 2*: Clinicopathological details of pnSTING scored cohort

| **Characteristics (N=156)** | **N** | **%** |
|---|---|---|
| ***Tumor Grade*** | | |
| **G1** | 1 | 0.6 |
| **G2** | 51 | 32.7 |
| **G3** | 104 | 66.7 |
| ***pN Stage (nodal involvement)*** | | |
| **N0** | 68 | 43.6 |
| **N1mi** | 2 | 1.3 |
| **N1** | 50 | 32.1 |
| **N3** | 21 | 23.5 |
| **N3** | 15 | 9.5 |
| ***pT stage*** | | |
| **T1** | 31 | 19.9 |
| **T2** | 93 | 59.6 |
| **T3** | 27 | 17.3 |
| **T4** | 5 | 3.2 |
| ***Histological Type*** | | |
| **Ductal (no special type)** | 128 | 82 |
| **Lobular (no special type)** | 14 | 9 |
| **Mixed ductal and lobular** | 11 | 7.1 |
| **Other** | 3 | 1.9 |
| ***ER status*** | | |
| **Positive** | 92 | 59 |
| **Negative** | 64 | 41 |

ER: Estrogen receptor; pT: pathological tumor stage; pN: pathological lymph node stage.

*Supplementary Table 3:* Relapse free survival stratified by pnSTING (low/high) in consensus breast subgroups

| | | N(n) | HR | 95% CI | p-value |
|---|---|---|---|---|---|
| ***Relapse Free Survival*** | | | | Univariate | |
| **ER+/HER2-/Ki67-** | low | 22 (5) | 1 | | |
| | high | 20 (0) | 0.1304[#] | 0.0022-0.7568 | 0.0232* |
| **ER+/HER2-/Ki67+** | low | 19 (7) | 1 | | |
| | high | 13 (2) | 0.3570 | 0.0959-1.329 | 0.1776 |
| **ER+/HER2+/Ki67+** | low | 9 (4) | 1 | | |
| | high | 9 (1) | 0.2347 | 0.0406-1.356 | 0.1574 |
| **ER-/HER2+** | low | 13 (5) | 1 | | |
| | high | 10 (2) | 0.4320 | 0.0981-1.903 | 0.3005 |
| **ER-/PR/HER2-** | low | 14 (3) | 1 | | |
| | high | 26 (7) | 1.327 | 0.3656-4.818 | 0.6805 |
| **ER+/HER2-/Ki67+/-** | low | 65 (23) | 1 | | |
| | high | 56 (8) | 0.3729 | 0.1843-0.7542 | 0.0121* |
| **ER+/-/HER2+** | low | 22(9) | 1 | | |
| | high | 19 (3) | 0.3196 | 0.1030-0.9912 | 0.0703 |
| **Non-ER-/PR-/HER2-** | low | 87 (32) | 1 | | |
| | high | 75 (11) | 0.3573 | 0.1965-0.6497 | 0.0020* |

[#]Mantel-Haenszel HR reported

*Supplementary Table 4:* pnSTING$^{low}$ gene signature

| |
|---|
| IMPA2 |
| SLC2A4 |
| CDT1 |
| CFAP47 |
| RABGGTB |
| NCAPD3 |
| ABRA |
| SLC35C2 |
| DNAH12 |
| SLC35E2B |
| LINC00479 |
| HYDIN2 |
| CSN3 |
| FAM71B |
| RCAN3 |
| MSH2 |
| PCDHA2 |
| SHISA3 |
| EXO1 |
| GALNT14 |
| VPS33A |
| DLGAP5 |
| HFM1 |
| BEND3 |
| MEX3B |

*Note: signature = -sum(gene expression)

*Supplementary Table 5:* pnSTING gene signature applied to independent datasets

| **Gene Signature** | | **N(n)** | **HR** | **%95 CI** | **p-value** |
|---|---|---|---|---|---|
| **Relapse Free Survival** | | | Univariate | | |
| **ER+ IHC cohort** | low | 46 (14) | 1 | | |
| | high | 46 (5) | 0.3220 | 0.1309-0.7924 | 0.0217* |
| **ER- IHC cohort** | low | 32 (9) | 1 | | |
| | high | 32 (8) | 0.8985 | 0.3472-2.325 | 0.8254 |
| **ER+ & ER- IHC cohort** | low | 78 (24) | 1 | | |
| | high | 78 (12) | 0.4402 | 0.2287-0.8471 | 0.0169* |
| **ER+ Non IHC cohort** | low | 38 (10) | 1 | | |
| | high | 38 (3) | 0.2795 | 0.09416-0.8295 | 0.0383* |
| **ER- Non IHC cohort** | low | 19 (10) | 1 | | |
| | high | 18 (7) | 0.6193 | 0.2389-1.605 | 0.3256 |
| **ER+ & ER- Non IHC cohort** | low | 58 (22) | 1 | | |
| | high | 58 (8) | 0.3126 | 0.1525-0.6409 | 0.0029** |
| **ER+ combined cohort** | low | 84 (24) | 1 | | |
| | high | 84 (8) | 0.3015 | 0.1506-0.6034 | 0.0018** |
| **ER- combined cohort** | low | 51(19) | 1 | | |
| | high | 50 (15) | 0.7740 | 0.3951-1.518 | 0.4565 |
| **ER+ & ER- combined cohort** | low | 135 (43) | 1 | | |
| | high | 135 (23) | 0.3773 | 0.2326-0.6121 | 0.0002*** |
| **METABRIC (2012&2016)** | | **N(n)** | **HR** | **%95 CI** | **p-value** |
| **Overall Survival** | | | Univariate | | |
| **ER+** | low | 486 (297) | 1 | | |
| | high | 485(262) | 0.8098 | 0.6855-0.9650 | 0.0125* |
| **ER-** | low | 167 (85) | 1 | | |
| | high | 166 (96) | 1.258 | 0.9394-1.684 | 0.1201 |
| **ER+ No chemo** | low | 433 (270) | 1 | | |
| | high | 432 (240) | 0.8322 | 0.6995-0.99 | 0.0379* |
| **ER+ Chemo** | low | 53 (29) | 1 | | |
| | high | 53 (20) | 0.5356 | 0.3046-0.9419 | 0.028* |
| **TCGA Nature 2012** | | **N(n)** | **HR** | **%95 CI** | **p-value** |
| **Relapse Free Survival** | | | Univariate | | |
| **ER+** | low | 189 (25) | 1 | | |
| | high | 191 (20) | 0.5481 | 0.3030-0.9953 | 0.0405* |
| **ER-** | low | 59 (10) | 1 | | |
| | high | 59 (7) | 1.441 | 0.5569-3.729 | 0.4251 |
| **GSE2034 (Wang et al)** | | **N(n)** | **HR** | **%95 CI** | **p-value** |
| **Relapse Free Survival** | | | Univariate | | |
| **ER+** | low | 104 (52) | 1 | | |
| | high | 105 (28) | 0.4292 | 0.2761-0.6672 | 0.0002*** |
| **ER-** | low | 39 (10) | 1 | | |
| | high | 38 (17) | 1.779 | 0.8364-3.783 | 0.1406 |

*Supplementary Table 6:* TP53, PIK3CA and MAP3K1 alterations in ER+ METABRIC and TCGA datasets stratified by pnSTING signature

| Dataset | Gene | No. cases with alteration (%) | | Log Ratio | p-value | q-value |
|---|---|---|---|---|---|---|
| **Mutations** | | STING Sig low | STING Sig high | | | |
| **META-BRIC** | TP53 | 179 (36.83) | 61 (12.58) | -1.55 | <10 e-10 | <10 e-10 |
| | PIK3CA | 185 (38.07) | 268 (55.26) | 0.54 | 5.26 e-8 | 4.549 e-6 |
| | MAP3K1 | 31 (6.38) | 83 (17.11) | 1.42 | 1.15 e-7 | 6.61 e-6 |
| **TCGA (Nature 2012)** | TP53 | 73 (38.02) | 19 (9.95) | -1.93 | 5.09 e-11 | 4.40 e-7 |
| | PIK3CA | 61 (31.77) | 94 (49.21) | 0.63 | 3.596 e-4 | 0.577 |
| | MAP3K1 | 11 (5.73) | 27 (14.14) | 1.3 | 4.55e-3 | 0.577 |

*Supplementary Table 7:* Copy number alterations of MYC and CCND1 in ER+ METABRIC and TCGA datasets stratified by pnSTING signature

| Dataset | Gene | No Cases with alteration (%) | | Log Ratio | p-value | q-value |
|---|---|---|---|---|---|---|
| **CNA** | | STING Sig low | STING Sig high | | | |
| **META-BRIC** | MYC (8q24,21) | 171 (35.19) | 77 (15.88) | -1.15 | <10 e-10 | 5.75 e-9 |
| | CCND1 (11q13.3) | 145 (29.63) | 56 (11.55) | -1.37 | <10 e-10 | 2.76 e-9 |
| **TCGA (Nature 2012)** | MYC (8q24,21) | 28 (14.81) | 16 (8.42) | -0.81 | 0.0369 | 0.563 |
| | CCND1 (11q13.3) | 50 (26.46) | 24 (12.63) | -1.07 | 5.035 e-4 | 0.166 |
